# A compact regulatory RNA element in mouse Hsp70 mRNA

**DOI:** 10.1101/2023.02.22.529618

**Authors:** Wenshuai Wang, Fei Liu, Maria Vera Ugalde, Anna Marie Pyle

**Affiliations:** Department of Molecular, Cellular and Developmental Biology, Yale University, New Haven, CT 06511, USA; Howard Hughes Medical Institute, Yale University, New Haven, CT 06520, USA; Department of Biochemistry. McGill University, Montreal, QuebecH3G 1Y6, Canada

## Abstract

Hsp70 performs molecular chaperone functions by assisting in folding newly synthesized or misfolded proteins, thereby counteracting various cell stresses and preventing multiple diseases including neurodegenerative disorders and cancer. It is well established that Hsp70 upregulation during post-heat shock stimulus is mediated by cap-dependent translation. However, the molecular mechanisms of Hsp70 expression during heat shock stimulus remains elusive, even though the 5’ end of Hsp70 mRNA may form a compact structure to positively regulate protein expression in the mode of cap-independent translation. The minimal truncation which can fold to a compact structure was mapped and its secondary structure was characterized by chemical probing. The predicted model revealed a highly compact structure with multiple stems. Including the stem where the canonical start codon is located, several stems were identified to be vital for RNA folding, thereby providing solid structural basis for future investigations on the function of this RNA structure on Hsp70 translation during heat shock.

## INTRODUCTION

Hsp70 (70 kilodalton heat shock protein) is an indispensable and highly conserved protein chaperone, which functions by stabilizing newly synthesized proteins to ensure correct protein folding and refolding proteins unfolded by cell stress, thereby counteracting the stress, modulating immune response and promoting cell survival^1–5^. The dysregulation of molecular chaperone Hsp70 has been associated with multiple diseases^5–7^. Therefore, understanding the mechanisms of Hsp70 translation regulation and gene expression is central to fully utilize the potential of Hsp70 for developing novel therapeutic strategies against diseases including neurodegenerative disorders.

Years of study has established that, upon stresses such as heat shock, the induced misfolded proteins interact with HSP (heat shock protein) to release transcription factor HSF1 (Heat shock factor 1), which is transported into nucleus to initiate the transcription of HSP^8,9^. As a result, HSP in the cytosol is dramatically upregulated, thereby efficiently activating molecular chaperone functions. However, compared with Hsp70 upregulation by cap-dependent translation during post-heat shock stimulus^10^, the translation mechanisms during heat shock stimulus are not well understood, especially when cap-dependent translation is prohibited^10^. This strongly suggests that Hsp70 can be expressed by elusive cap-independent translation mechanisms under stress conditions. It is well known that certain RNA structures including IRES motifs (Internal Ribosome Entry Site), located in the 5’ end of mRNA, can help recruit ribosomes to initiate cap-independent translation^10^. As the 5’ UTR of Hsp70 mRNA has been reported to play positive roles during cap-independent Hsp70 translation^11–16^, we hypothesized that the 5’ end of Hsp70 mRNA may fold into a compact structure that assists in cap-independent Hsp70 translation during heat shock stress.

To determine the compact structure formed by the 5’ end of Hsp70 mRNA, we examined the influence of of Hsp70 mRNA truncations on RNA folding, and then we probed the RNA secondary structure and identified unique features that contribute to formation of the compact structure. To our surprise, we found that the minimal construct (UTR26) that can fold to a compact structure is composed of 5’ UTR and part of the ORF sequence of Hsp70 mRNA. UTR26 contains a network of stable RNA stems, including a stem that contains the start codon. These motifs are vital for RNA folding, thereby providing a structural basis for understanding the function of a putative regulatory RNA structure on Hsp70 translation during heat shock.

## RESULTS

### A compact structure in 5’ end of Hsp70 mRNA

We hypothesized that the 5’ end of Hsp70 mRNA folds to a compact structure, potentially acting as a thermosensor to mediate cap-independent translation of Hsp70 during heat shock. To test this hypothesis, we *in-vitro* transcribed the 5’ end of Hsp70 mRNA and used native gel electrophoresis to test its ability to fold into a compact state under normal (37 °C) and heat shock (43 °C) conditions, where the folded RNAs migrate faster in a gel than unfolded RNAs. Surprisingly, we found that the 5’ UTR (untranslated region, UTR1) in isolation cannot fold to a compact structure *in vitro*, supported by the lack of folded RNA band for the UTR-only construct (Figure 1A and 1B). In comparison, the addition of 100 nucleotides of ORF (open reading frame) sequence to the UTR (resulting in construct UTR7) enables the RNA to fold to a compact structure under both normal and heat shock conditions (Figure 1A and 1B), suggesting an important role for ORF sequence in this functional element. However, with 30-nt, 54-nt, 84-nt and 150-nt ORF additions (UTR2, UTR4, UTR10 and UTR9), the RNA constructs did not fold better than UTR7 (Figure 1A, 1B and 1C), confirming the unique importance of the 100-nt ORF in UTR7 in the compact structure. To explore the minimal sequence required for the folding process, we fine-tuned the ORF length based on UTR7 and tested folding of the resultant constructs. Intriguingly, when compared with UTR7, the removal of 4 nts from the ORF of UTR7 (resulting in UTR26) achieved better folding, and a temperature-dependence to folding, as illustrated by the more homogenous and temperature-sensitive folded RNA band in the native gel (Figure 1D and 1E). And all other RNA truncations failed to enhance folding (Figure 1D), suggesting that the 3’ end of UTR7 and UTR26 is heavily involved in the compaction process and that it may contain important secondary structures. These findings suggest a model involving compact structure in the 5’ end of Hsp70 mRNA and we have established an identified minimal RNA construct for further characterization of component secondary structures.

**Figure 1.**
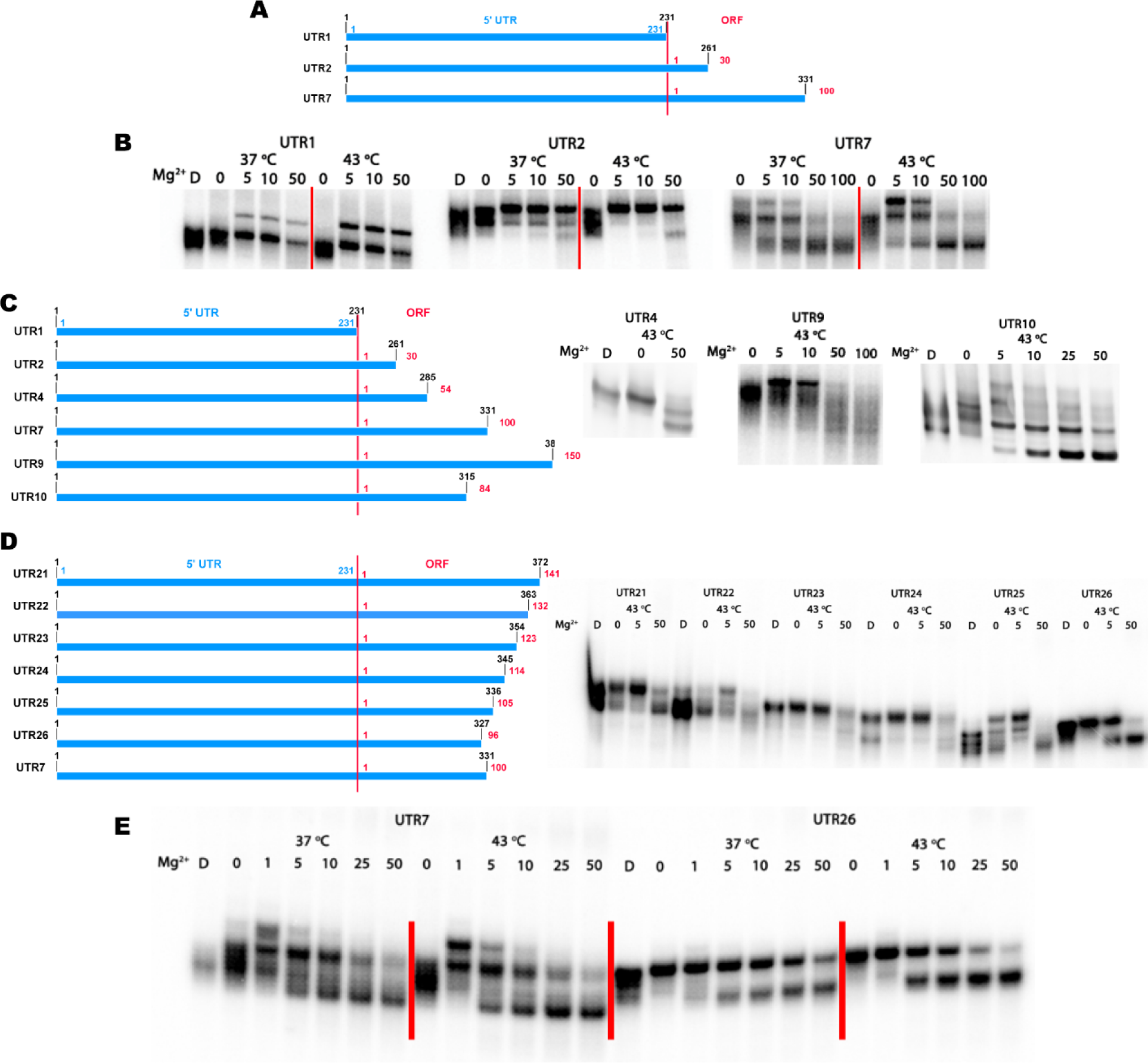
The 5’ end of Hsp70 mRNA folds to a compact structure. (A) Schematic representation of different RNA constructs containing 5’ end of Hsp70 mRNA. The 5’ UTR and ORF sequences are denoted in blue and red, respectively. (B) Folding results of RNA constructs described above in native gel at 37 °C and 43 °C. The denatured RNA sample is labeled as “D”. The Mg^2+^ concentrations are shown as 0, 5, 10, 50 mM. (C-D) Schematic representation of different RNA constructs for determining the minimal folding truncation, and corresponding folding results of these truncations. (E) Folding results of two best truncations (UTR7, UTR26) showing well defined folded RNA band.

### Determination of the secondary structure

Knowing that the minimal construct UTR26 folds to a compact structure, we utilized SHAPE (Selective 2’ -hydroxyl acylation analyzed by primer extension) and DMS probing to characterize its secondary structure. Having separated a well-folded RNA using a size-exclusion column (see methods, Figure S1), the secondary structure of UTR26 RNA at 43 °C was examined by using SHAPE and DMS probing. The SHAPE reagent 1-methyl-7-nitroisatoic anhydride (1M7) selectively acetylates the 2’ -hydroxyl group of RNA nucleotides with flexible backbones, and DMS selectively methylates the heterocyclic nitrogen atoms on unbasepaired adenines and cytosines^17,18^. A robust secondary structure is established by the agreement between these two canonical chemical probing approaches. We monitored the SHAPE reactivity at single-nucleotide resolution and used normalized SHAPE reactivities as pseudo-free energy constraints to restrain the secondary structure predication by minimum free energy (MFE, Figure 2A). To evaluate the SHAPE-restrained secondary structure model, we then aligned the DMS reactivity to the model, observing that the DMS reactivity data matched well with the SHAPE reactivity (Figure 2A), suggesting good agreement between the two methods. The predicted model is mainly composed of eight stem loops (H1-H8, Figure 2A), indicating that UTR26 RNA is a highly compact structure with about 55% of the nucleotides base-paired (Figure 2A). Interestingly, both 5’ and 3’ ends of UTR26 fold to stem-loop elements, emphasizing their importance during the folding process. More importantly, the canonical star codon comprises stem H6, suggesting a potential role of stem H6 on Hsp70 translation during heat shock.

**Figure 2.**
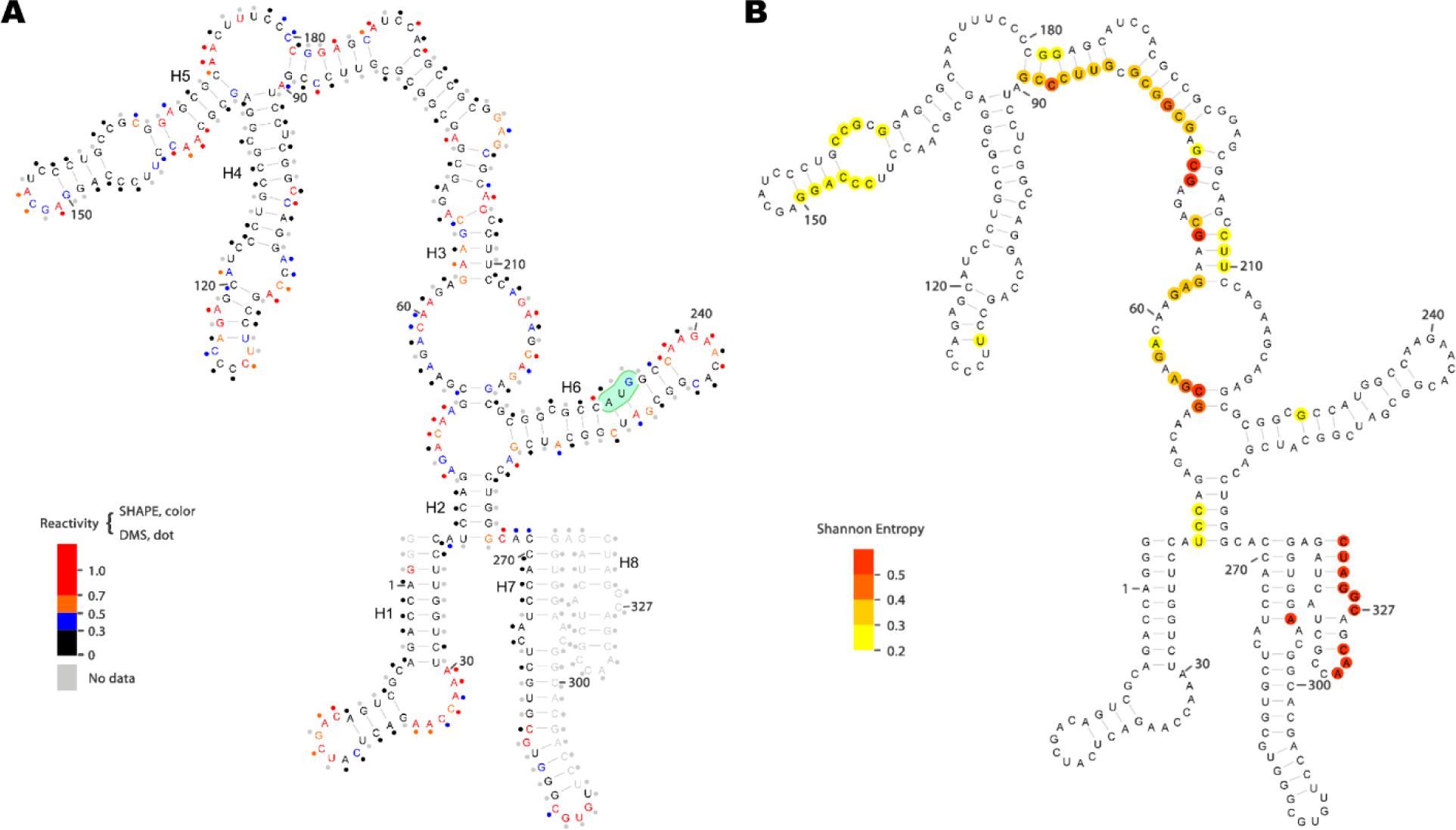
Predicted secondary structure of 5’ end of Hsp70 mRNA. (A) SHAPE reactivities are presented by colored nucleotides, while DMS reactivities are rendered as colored dots next to the nucleotides. Highly reactive nucleotides are colored in red or orange. Nucleotides with low reactivities are shown in black or blue. Nucleotides with no data are in gray. The canonical start codon, AUG, is highlighted in green. (B) Nucleotides with high Shannon Entropy values are colored in red or orange. Nucleotides with medium Shannon Entropy values are rendered in dark or light yellow. Nucleotides with low Shannon Entropy values are not highlighted in the map.

### Validation of the compact structure

Having obtained a preliminary secondary structure model of UTR26 RNA, we next evaluated this model by testing a series of unzipping and compensatory mutants. The unzipping mutants were designed to disrupt folding to the compact state, while the corresponding compensatory mutants were designed to restore disrupted folding capacity. To determine the stem loops to be unzipped, we evaluated the potential local structural heterogeneity by calculating the Shannon entropy of each nucleotide based on the SHAPE-directed base-pair probabilities (Figure 2B). The average Shannon entropy was about 0.1, indicating an overall structural homogeneity of the UTR26 conformation. Among all eight stem loops, nucleotides of stem loop H1, H4 and H6 had much lower Shannon entropy (< 0.2), suggesting that these three stem loops are more stable (Figure 2B). We then introduced broad unzipping mutations to the three stem loops H1, H4 and H6, respectively, and tested the relative compaction of the unzipping mutants as a metric for stable folding. Based on the observed electrophoretic mobilities, the folding capacity of all the unzipping mutants, which destabilize H1, H4 and H6, was lost or attenuated, even at 50 mM Mg^2+^ (Figure 3A, 3B and 3C). To further evaluate the importance of specific stems on UTR26 folding, we introduced the corresponding compensatory mutations to the unzipping mutants, thereby restoring folding capability. By utilizing the same method mentioned above, we found that the compensatory mutants restored the folding that had been disrupted by the unzipping mutations (Figure 3A, 3B and 3C), confirming the predicted structural model and revealing that specific RNA stem motifs are important for RNA folding and for supporting the architecture of the RNA motif that regulates Hsp70 translation during heat shock.

**Figure 3.**
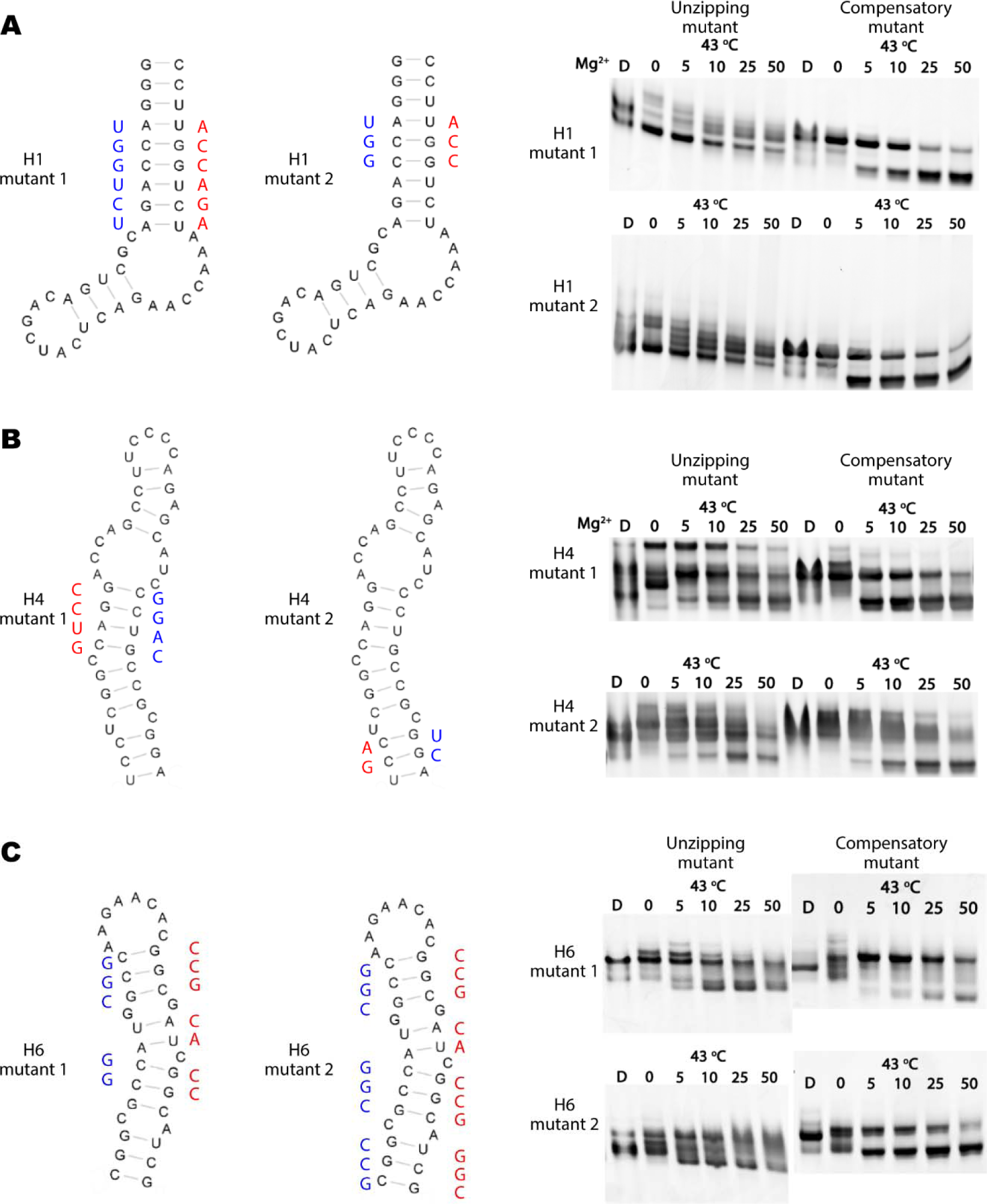
Validation of the predicted secondary structure. (A-C) Design of unzipping and compensatory mutants based on H1 (A), H4 (B) and H6 (C) stems. The unzipping mutants are in red. Based on the unzipping mutations, the corresponding compensatory mutants are presented by the red and blue nucleotides. The folding results of these mutants were displayed as described in Figure 1.

## DISCUSSION

Using *in-vitro* transcribed and folded RNA, we have successfully characterized the secondary structure at the 5’ end of Hsp70 mRNA, showing that it folds into a compact structure composed of multiple stems. To validate this *in vitro* result, it will be important to monitor the secondary structure of endogenous Hsp70 mRNA under normal, heat shock and post-heat shock conditions using *in vivo* chemical probing technique, such as *in vivo* SHAPE-MaP^19,20^. More insights of the structure on its function will be revealed by a comprehensive comparison between the *in vitro* and *in vivo* predicted models.

Although these results suggest the existence of a thermally-sensitive, compact structure in upstream regions of Hsp70 mRNA, exploring a functional role for this structure will require a cell-based assay for linking RNA structure in this region with Hsp70 expression during heat shock. In previous studies, a bicistronic assay has been used to study the effect of RNA sequence on protein translation, as in studies of IRES elements^10^. However, bicistronic assays of Hsp70 function have yielded conflicting results. In some cases they have indicated that the 5’ UTR of Hsp70 mRNA performs IRES or enhancer functions^14,15,21^, while in other cases, IRES activity was contraindicated^16,22^. Such opposing results on 5’ UTR of Hsp70 mRNA suggest that traditional bicistronic assays may not be suitable for exploring the influence of Hsp70 mRNA structure on its translation. For example, the ORF sequence encoded by UTR26 contains several amino acid residues that might influence subsequent translation or folding of the adjacent UTR26-fused reporter gene, thereby affecting the bicistronic assay results. Future studies would benefit from the use of gene editing to create a series of Hsp70 mutant cell lines that enable one directly monitor RNA structure *in-vivo* and the corresponding response of Hsp70 expression to heat shock.

It is likely that post-transcriptional modifications on Hsp70 mRNA contribute to the folding transition that we report here. RNA modifications have been shown to influence the conformational equilibria of RNA motifs and previous studies have shown that m(6)A modifications on Hsp70 mRNA can regulate cap-independent Hsp70 translation^11–13^. The A103 of Hsp70 mRNA was identified as a key m(6)A modification site, as A103C mutation decreases Hsp70 translation dramatically^13^. Interestingly, A103 has low SHAPE reactivity and may base pair with U122, which also has low SHAPE reactivity, despite the fact that these nucleotides were not predicted as base-paired in stem H4 (Figure 2A). This suggests the existence of more than one conformation in that position and that disruption of A103:U122 may break the balance in stability of an important motif, destabilizing the whole compact structure. As the predicted model in this study is based on data from the study of unmodified RNA transcripts, it is still an open question how RNA modifications, including m(6)A, modulate riboregulatory motifs and regulate Hsp70 translation.

## AUTHOR CONTRIBUTIONS

A.M.P., W.W. and F.L. designed the experiments. W.W. and F.L. performed the experiments and analyzed the data. M.V.U. helped generate ideas and provided plasmids. A.M.P. and W.W. wrote the paper.

## ACKNOWLEDGMENTS

We thank Robert Singer for helpful discussions. This work was supported by HHMI and by NIH Grant R01AI131518.

---

**Figure S1.**
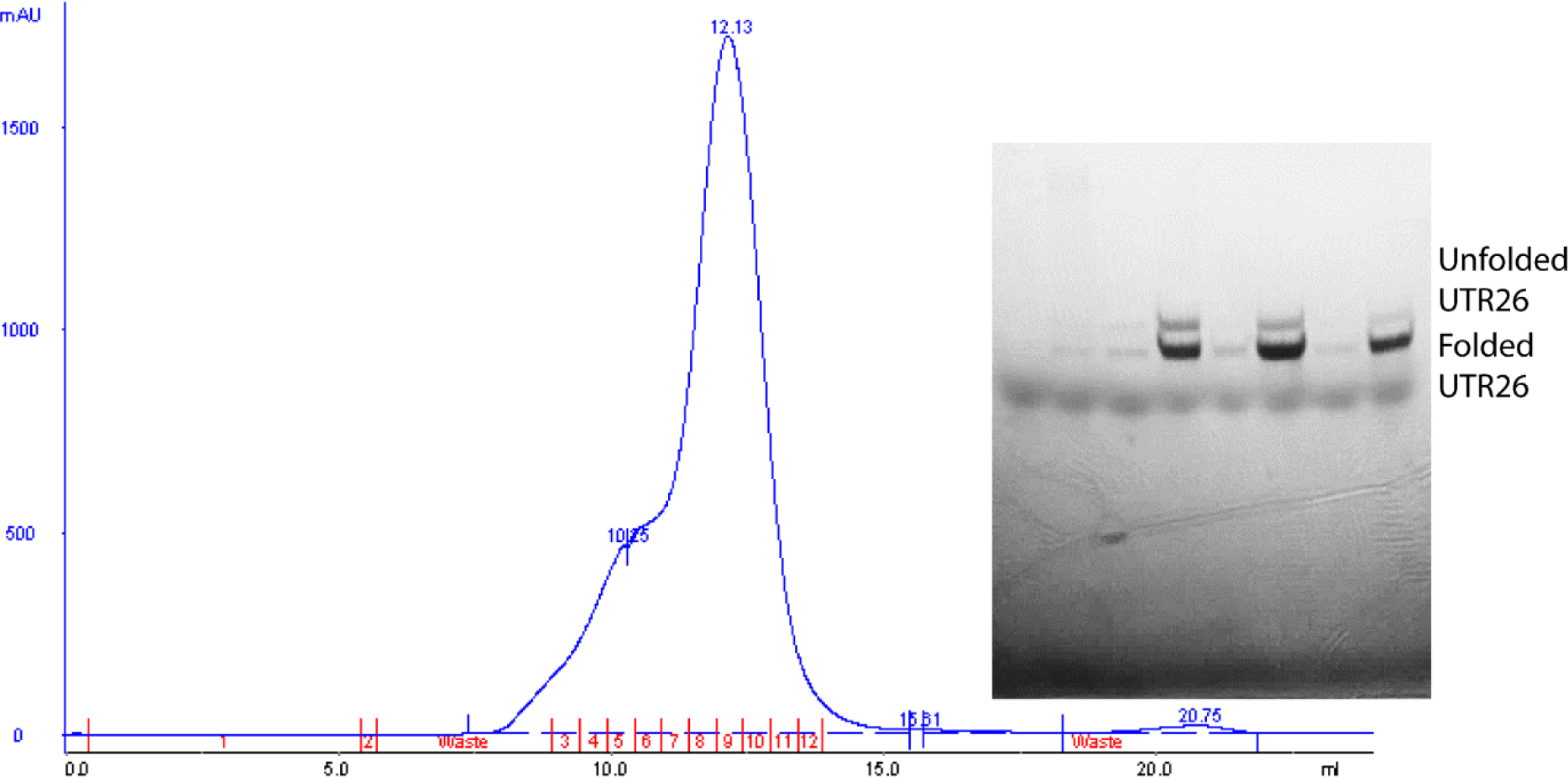
Large scale purification of folded UTR26. The elution volume of UTR26 after folding from SEC column (24 ml) is rendered. The fraction peak of eluted monomeric folded UTR26 is around 12.13 ml. The eluted UTR26 was assessed using native gel, and folded UTR26 was well separated from unfolded UTR26, especially in the fractions from the tail of the peak. This confirms that the RNA sample used for chemical probing is mainly composed of folded UTR26.

## MATERIALS AND METHODS

### Plasmids

The pBluescript vector (Agilent) was used for RNA expression *in vitro*. A series of DNA constructs being transcribed as the 5’-end of Hsp70 mRNA were cloned into the plasmid. Truncations and mutants were generated using Q5 Site-Directed Mutagenesis Kit (NEB).

### *In vitro* transcription and denatured purification of RNA

The RNA was *in-vitro* transcribed using T7 RNA polymerase with linearized plasmids as template. A 300 μl transcription solution contained 30 μg of linearized plasmids, 40 mM Tris-HCl (pH 8.0), 22 mM MgCl2, 10 mM DTT, 2 mM spermidine, 0.01% Triton X-100, 4 mM of each NTPs (ATP, UTP, CTP), 5 mM of GTP, 80 U of RNaseOUT™ Recombinant Ribonuclease Inhibitor (Thermo Fisher), and 25 μl of T7 RNA polymerase (lab made). With 2-hour incubation at 37 °C, the transcribed p3SLR30 was purified by gel extraction from 5% urea denaturing polyacrylamide gel.

### Native folding test

1 μM denatured RNA was folded in folding buffer containing 20 mM HEPES-K (pH 7.5), 0.1 mM EDTA-Na (pH 8.0), 50 mM KCl and appropriate MgCl_2_ concentrations. With 40-min incubation at 43 °C, the RNA folding state was monitored by running 5% polyacrylamide native gel.

### Chemical probing

The RNA construct namely UTR26 was folded as described above, and then purified by size-exclusion chromatography using S200 10/300 column (GE) equilibrated in buffer containing 20 mM Potassium Cacodylate (pH 7.0), 0.1 mM EDTA-Na (pH 8.0), 50 mM KCl, 5 mM MgCl_2_.

#### Selective 2’ -hydroxyl acylation analyzed by primer extension

The positive reaction was performed with 5 pmol of folded UTR26 incubated with 5 mM 1M7 (1-methyl-7-nitroisatoic anhydride)^17^, which was freshly dissolved in anhydrous DMSO, at 37 °C for 10 min, and the negative reaction was initiated with pure anhydrous DMSO. Reactions were quenched with buffer containing 1.5 M NaOAc (pH 5.2), 0.25 M EDTA-Na (pH 8.0) and 0.5 mg/ml glycogen (Thermo Fisher). Following precipitation with 70% ethanol, the precipitated RNA sample was dissolved in ddH_2_O and cleaned using RNA clean & concentrator kit (Zymo Research). Reverse transcription (RT) was carried out as previously described^23^. In brief, 0.1 μM 6-JOE-conjugated reverse primer (5’-6-JOE-GATGATCTCCACCTTGCCGT-3’) was extended with 1 pmol of RNA as template using Superscript III (Thermo Fisher) as reverse transcriptase. The quenching and DNA precipitation were performed as described above.

#### DMS probing

The reaction was performed with 5 pmol of folded UTR26 incubated with 0.077% v/v DMS (dimethyl sulfate, Sigma-Aldrich) dissolved in 100% ethanol at 37 °C for 10 min, and the negative reaction was initiated with pure ethanol. The reaction was quenched with 5% 2-Mercaptoethanol in ethanol. The sample preparation and RT were carried out the same as described above.

### Sequencing and structure mapping by capillary electrophoresis

The synthesis of fluorescent primers, preparation of sequencing ladders and capillary electrophoresis were performed as previously described^23^.

#### Data processing, normalization, and error assessment

Both SHAPE and DMS data were processed with QuShape as previously described^23,24^. The SHAPE reactivity profiles for each nucleotide were obtained by subtracting the peak areas of negative reaction (-) from the peak areas of the positive reactions (+), followed by normalization^24^. The DMS reactivity of adenosines and cytosines were normalized separately due to the intrinsic different reaction rates of DMS^18^. All the SHAPE and DMS probing were reproduced three times.

#### Structure determination

The software RNAStructure (https://rna.urmc.rochester.edu/RNAstructure.html) was used to predict the secondary structure of UTR26 with the SHAPE reactivity as pseudo-energy constraints^25^. The resulting predicted structures were manually assessed using the DMS data. The secondary structure was drawn with VARNA (http://varna-gui.software.informer.com/)

#### Shannon entropy calculation

The Shannon entropy was calculated as previously described^26,27^.

### Data availability

All chemical probing data are available upon request.

## REFERENCES

1 Molto, M. D., Pascual, L. & de Frutos, R. Puff activity after heat shock in two species of the Drosophila obscura group. Experientia 43, 1225–1227 (1987). https://doi.org:10.1007/BF01945535

2 Craig, E. A. & Gross, C. A. Is hsp70 the cellular thermometer? Trends Biochem Sci 16, 135–140 (1991). https://doi.org:10.1016/0968-0004(91)90055-z

3 Radons, J. The human HSP70 family of chaperones: where do we stand? Cell Stress Chaperones 21, 379–404 (2016). https://doi.org:10.1007/s12192-016-0676-6

4 Saibil, H. Chaperone machines for protein folding, unfolding and disaggregation. Nat Rev Mol Cell Biol 14, 630–642 (2013). https://doi.org:10.1038/nrm3658

5 Sherman, M. Y. & Gabai, V. L. Hsp70 in cancer: back to the future. Oncogene 34, 4153–4161 (2015). https://doi.org:10.1038/onc.2014.349

6 Joshi, S. et al. Adapting to stress - chaperome networks in cancer. Nat Rev Cancer 18, 562–575 (2018). https://doi.org:10.1038/s41568-018-0020-9

7 Muchowski, P. J. & Wacker, J. L. Modulation of neurodegeneration by molecular chaperones. Nat Rev Neurosci 6, 11–22 (2005). https://doi.org:10.1038/nrn1587

8 Vabulas, R. M., Raychaudhuri, S., Hayer-Hartl, M. & Hartl, F. U. Protein folding in the cytoplasm and the heat shock response. Cold Spring Harb Perspect Biol 2, a004390 (2010). https://doi.org:10.1101/cshperspect.a004390

9 Gomez-Pastor, R., Burchfiel, E. T. & Thiele, D. J. Regulation of heat shock transcription factors and their roles in physiology and disease. Nat Rev Mol Cell Biol 19, 4–19 (2018). https://doi.org:10.1038/nrm.2017.73

10 Leppek, K., Das, R. & Barna, M. Functional 5’ UTR mRNA structures in eukaryotic translation regulation and how to find them. Nat Rev Mol Cell Biol 19, 158–174 (2018). https://doi.org:10.1038/nrm.2017.103

11 Meyer, K. D. et al. 5’ UTR m(6)A Promotes Cap-Independent Translation. Cell 163, 999–1010 (2015). https://doi.org:10.1016/j.cell.2015.10.012

12 Coots, R. A. et al. m(6)A Facilitates elF4F-lndependent mRNA Translation. Mol Cell 68, 504–514 e5O7 (2017). https://doi.org:10.1016/i.molcel.2017.10.002

13 Zhou, J. et al. Dynamic m(6)A mRNA methylation directs translational control of heat shock response. Nature 526, 591–594 (2015). https://doi.org:10.1038/nature15377

14 Rubtsova, M. P. et al. Distinctive properties of the 5’-untranslated region of human hsp70 mRNA. J Biol Chem 278, 22350–22356 (2003). https://doi.org:10.1074/jbc.M303213200

15 Vivinus, S. et al. An element within the 5’ untranslated region of human Hsp70 mRNA which acts as a general enhancer of mRNA translation. Eur J Biochem 268, 1908–1917 (2001). https://doi.org:10.1046/j.1432-1327.2001.02064.x

16 Sun, J., Conn, C. S., Han, Y., Yeung, V. & Qian, S. B. PI3K-mTORCl attenuates stress response by inhibiting cap-independent Hsp70 translation. J Biol Chem 286, 6791–6800 (2011). https://doi.org:10.1074/jbc.M110.172882

17 Mortimer, S. A. & Weeks, K. M. A fast-acting reagent for accurate analysis of RNA secondary and tertiary structure by SHAPE chemistry. J Am Chem Soc 129, 4144–4145 (2007). https://doi.org:10.1021/ja0704028

18 Rouskin, S., Zubradt, M., Washietl, S., Kellis, M. & Weissman, J. S. Genome-wide probing of RNA structure reveals active unfolding of mRNA structures in vivo. Nature 505, 701–705 (2014). https://doi.org:10.1038/nature12894

19 Siegfried, N. A., Busan, S., Rice, G. M., Nelson, J. A. & Weeks, K. M. RNA motif discovery by SHAPE and mutational profiling (SHAPE-MaP). Nat Methods 11, 959–965 (2014). https://doi.org:10.1038/nmeth.3029

20 Huston, N. C. et al. Comprehensive in vivo secondary structure of the SARS-CoV-2 genome reveals novel regulatory motifs and mechanisms. Mol Cell 81, 584–598 e585 (2021). https://doi.org:10.1016/j.molcel.2020.12.041

21 Rocchi, L., Alfieri, R. R., Petronini, P. G., Montanaro, L. & Brigotti, M. 5’-Untranslated region of heat shock protein 70 mRNA drives translation under hypertonic conditions. Biochem Biophys Res Commun 431, 321–325 (2013). https://doi.org:10.1016/j.bbrc.2012.12.100

22 Andreev, D. E. et al. Differential contribution of the m7G-cap to the 5’ end-dependent translation initiation of mammalian mRNAs. Nucleic Acids Res 37, 6135–6147 (2009). https://doi.org:10.1093/nar/gkp665

23 Somarowthu, S. et al. HOTAIR forms an intricate and modular secondary structure. Mol Cell 58, 353–361 (2015). https://doi.org:10.1016/j.molcel.2015.03.006

24 Karabiber, F., McGinnis, J. L., Favorov, O. V. & Weeks, K. M. QuShape: rapid, accurate, and best-practices quantification of nucleic acid probing information, resolved by capillary electrophoresis. RNA 19, 63–73 (2013). https://doi.org:10.1261/rna.036327.112

25 Low, J. T. & Weeks, K. M. SHAPE-directed RNA secondary structure prediction. Methods 52, 150–158 (2010). https://doi.org:10.1016/i.ymeth.2010.06.007

26 Liu, F., Somarowthu, S. & Pyle, A. M. Visualizing the secondary and tertiary architectural domains of IncRNA RepA. Nat Chem Biol 13, 282–289 (2017). https://doi.org:10.1038/nchembio.2272

27 Mathews, D. H. Using an RNA secondary structure partition function to determine confidence in base pairs predicted by free energy minimization. RNA 10, 1178–1190 (2004). https://doi.org:10.1261/rna.7650904

